# Dispersers are more likely to follow mucus trails in the land snail *Cornu aspersum*

**DOI:** 10.1101/373001

**Authors:** Alexandre Vong, Armelle Ansart, Maxime Dahirel

## Abstract

Dispersal, i.e. movement leading to gene flow, is a fundamental although costly life history trait. The use of indirect social information may help mitigate these costs, yet in many cases little is known about the proximate sources of such information, and how dispersers and residents may differ in their information use. Land gastropods, which have a high cost of movement and obligatorily leave information potentially exploitable by conspecifics during movement (through mucus trails), are a good model to investigate links between dispersal costs and information use. We used Y-mazes to see whether dispersers and residents differed in their trail-following propensity, in the snail *Cornu aspersum*. Dispersers followed mucus trails more frequently than expected by chance, contrary to non-dispersers. Ignoring dispersal status during tests would lead to falsely conclude to no trail-following for the majority of ecologically realistic scenarios. Trail following by dispersers may reduce dispersal costs by reducing energy expenditure and helping snails find existing patches. Finally, we point that ignoring the potential for collective dispersal provided by trail-following abilities may lead to wrong inferences on the demographic and genetic consequences of dispersal.

## Introduction

Dispersal, i.e. movement potentially leading to gene flow in space, is a key life history trait connecting ecological and evolutionary dynamics [1–3]. Costs and uncertainty associated with dispersal [2] can be reduced by gathering information about current and prospective habitats [4–6]. Indirect social information, obtained from the presence, density, traits and/ or performance of conspecifics, although less accurate than direct experience, may provide information about potential nearby habitats without the need for costly prospecting [4,5,7].

Movement in terrestrial gastropods (snails and slugs) is among the costliest in animals, due to mucus secretion leading to substantial energy and water loss even over short distances [8,9]. As mucus production is obligatory, many crawling gastropods have unsurprisingly evolved trail following behaviour to locate conspecifics, mates, or potential gastropod prey [10]. Information on phenotype (which can inform on origin patch quality [5]) can additionally be gathered from mucus trail physical and chemical characteristics [10]. Crawling on pre-existing trails may also reduce the need for mucus production, leading to significant energy savings [11]. However, trail following has actually mostly been studied in aquatic gastropods, and knowledge about its frequency and function in terrestrial snails is limited [10].

The brown garden snail *Cornu aspersum* (Müller) (Helicidae; syn. *Helix aspersa*) is a well-studied generalist land snail, able to thrive and disperse even in strongly fragmented, urban habitats [12,13]. *Cornu aspersum* snails are sensitive to mucus accumulations [14] and adjust their emigration and immigration decisions to conspecific density [15]. They appear to follow trails slightly more than expected by chance [16], but there is no clear evidence so far that they use trails during dispersal or that dispersers and residents react differently to social information. Using a Y-maze setup, we here tested the hypotheses that *Cornu aspersum* snails are trail followers, and that dispersers would be more likely to follow trails; indeed, they would benefit more from potential energy savings and from the social information about conspecific presence than residents, which are not expected to stray far from an already established group of conspecifics.

## Methods

### Rearing conditions

Adult and older subadult brown garden snails (greater shell diameter > 25 mm) were obtained from two sources in April - May 2016. First, we selected 50 individuals (see below for details) from a set of snails used in a previous dispersal study [17] and collected from natural populations in parks in Rennes, France (≈1°38’W, ≈48°7’N, hereafter the “natural population”). We also tested 47 individuals randomly chosen from a set of 130 snails obtained from a snail farm in Corps-Nuds, close to Rennes (1°36’37″ W, 47°58’44″N, hereafter the “farm population”). Snails were kept under controlled conditions (20 ± 1 C; 16L: 8D; *ad lib* cereal-based snail food, Hélinove, Le Boupère, France), in polyethylene boxes covered by a net (30 × 45 × 8 cm) and lined by synthetic foam kept saturated with water. They were housed in groups of at most forty before use, and then in groups of eight to ten individually marked snails (with felt-tip paint markers) for at least one week before dispersal tests. Boxes were cleaned and linings changed once per week.

### Dispersal tests

We assessed dispersal in an outdoor asphalted area on the Beaulieu university campus, Rennes (1°38’15″O, 48°6’59″N; see [17] for methods details and their relevance to *Cornu aspersum* ecology). Briefly, rearing boxes (including food and water) were placed in the middle of the test area and left open for one night (19:00 to 09:00). Snails found more than 1 m outside of the box in the morning, i.e. beyond the typical home range, were considered dispersers. In the natural population, a random set of 25 dispersers and 25 residents was then selected for further experiments. For logistical reasons, trail following was tested before dispersal tests in the farm population; such a balanced design could not therefore be attained (15 out of 47 tested snails from that population were dispersers).

### Trail following experiment

We tested snail trail following propensity using Y-mazes [10] in a dark room, as snails are nocturnal (Fig. 1). The experimenter (AV) wore latex gloves during setup and experiments to limit uncontrolled disturbance by human odours. Mazes were lined with watered synthetic foam (as in rearing boxes), and 7g of snail food were placed at the extremities of both choice arms to stimulate movement. To limit escapes, Y-mazes were raised by 11 cm and the stand on which the starting (main) arm of the maze rested was covered in soot, repulsive to snails ([18], Fig. 1). First, a “marker” snail randomly chosen among untested stock snails was placed in the main arm of the maze and left free to move for 10 minutes. Marker snails that made U-turns or used both choice arms were excluded from further tests. Within 10 minutes after removing the marker, a “tracker” snail was then placed at the start of the maze and left free to move for 10 minutes. Tracker snails were counted as trail followers if they chose the same arm as the marker snail. New maze linings and feeders were used for each test. Before experiments, preliminary tests with no marker snail were done to confirm that there was no intrinsic left-right bias in choices [10] (47.5 % chose the left side, binomial test against a 50% expectation, N = 40, p = 0.87). Following this, left-right symmetry during actual tests was enforced by alternately proposing left-side trails and right-side trails to successive tracker snails, randomly selected from simultaneously generated trails.

**Figure 1.**
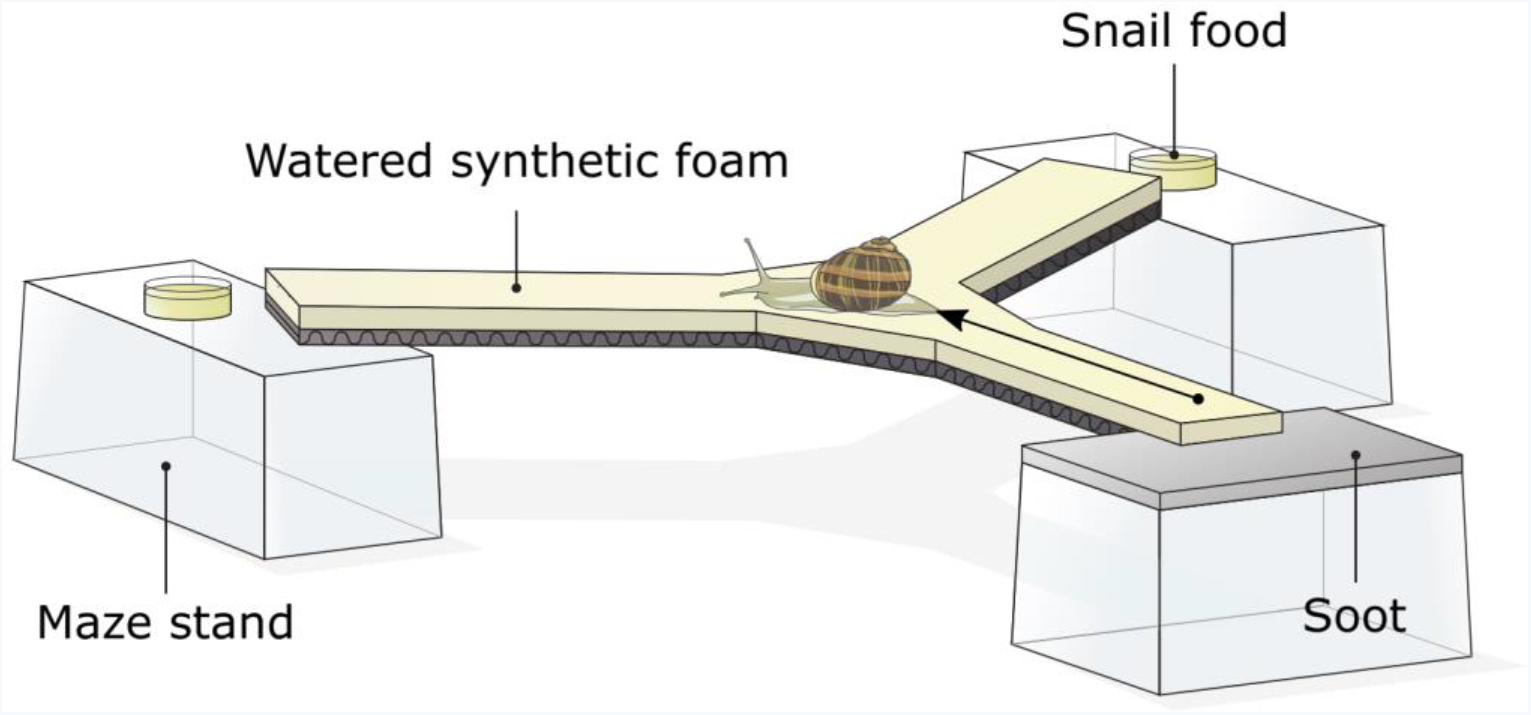
Experimental setup (Y-maze) for the trail-following experiments (not to scale; arm width: 3.5 cm; main arm length: 10 cm; choice arm length: 15 cm; angle between arms: 120°).

### Statistical analyses

We used a binomial generalized linear model to test for an effect of dispersal status, population of origin and their interaction on trail following probability. Analyses were done using R, version 3.5.1 [19], and the *car* and *emmeans* packages [20,21].

## Results

Dispersers were more likely to follow trails than residents (*Χ^2^_1_* = 6.40, *p* = 0.01, Fig. 2). Contrary to dispersers, residents were not more likely to follow trails than the 50% expected by chance (see 95% confidence intervals on Fig. 2). There was no significant effects of population of origin or dispersal status × population interaction (*Χ^2^_1_* = 0.17 and 1.20, *p* = 0.68 and 0.27, respectively).

**Figure 2.**
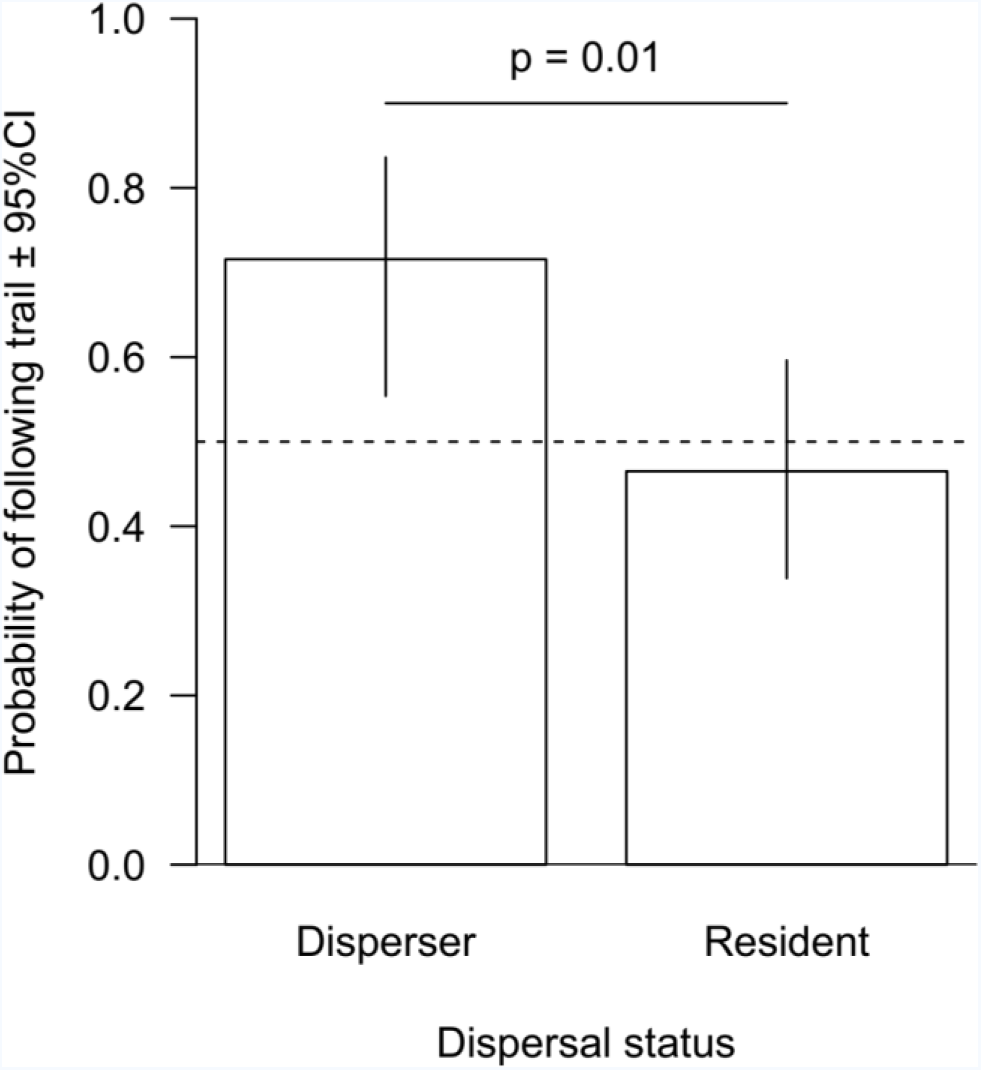
Trail following rate as a function of dispersal status (model predictions and 95% confidence intervals based on binomial GLM, the non-significant effect of origin population is averaged out; N = 97).

## Discussion

Dispersers, but not residents, were more likely to follow trails than expected by chance, indicating that mucus trails are usable sources of indirect social information in *Cornu aspersum* snails. A non-exclusive alternative is that trail-following is an energy saving measure [11], which would be more useful for dispersers. Intuitively and importantly, our results also indicate that tests realized without knowledge of dispersal status may falsely conclude to the absence of trail following behaviour under ecologically realistic dispersal rates (see Supplementary Material).

Mucus trails may even have higher value for dispersers compared to previously studied sources of indirect social information, as they not only give information about meta-population level habitat quality or population density [4,5,7], but also about the spatial location of other patches (or at least other snails), further reducing dispersal costs. This may be especially valuable in highly fragmented urban areas, where artificial porous substrates may make movement more costly [9] and inter-patch distances are often larger than the (low) perceptual range of *C. aspersum* [12]. Note that due to our experimental design, we here only confirm that snails follow trails with positive polarity, i.e. by going in the same direction as the trail layer. In this context, a way to use trails as cues of prospective habitat location would be to follow snails homing back to their roosts [16]. If they are able to follow trails with negative polarity (which is not unlikely [10]), they might additionally be able to “walk back” trails left by immigrants to reach their departure point.

The well-documented effects of within-habitat mucus accumulations on life-history and behaviour are size and species-specific (e.g. [14]), and recent evidence suggest this is also the case for trail following in at least one land snail group [22]. An important next step will be to determine how social information and phenotype combine to shape dispersal, especially in the context of matching habitat choice [3]. Furthermore, dispersers following trails (potentially laid by previous dispersers) may also provide a mechanism for collective dispersal in snails, several individuals following an initial trail-blazer [23]. As pointed out by Cote et al. [23], such collective dispersal, whether involving kin or non-kin, would have wide-ranging yet poorly studied consequences for population dynamics, evolution and genetic structure, and affect our ability to infer spatial dynamics from population genetics data. Land snails, by combining ease of behavioural study in controlled and naturalistic conditions, trail following ability and a long and ongoing history as population genetic models [13,24], may be one of the best study taxa yet to investigate these questions.

## Acknowledgements

We warmly thank Baptiste Averly for his involvement in the preliminary experiments that led to the experimental design presented in this study.

## Author contributions (in alphabetical order within each category)

Original idea: AA, MD; snail collection: AA, AV, MD; study design: AA, AV, MD; data collection: AV; data analysis: AV, MD; manuscript initial draft: MD, from an initial report by AV. All authors contributed critically to the draft and gave final approval for publication.

## Data accessibility

Data are available on Figshare (doi: 10.6084/m9.figshare.6840179)

## Funding

MD was funded by a Fyssen Foundation postdoctoral grant.

## Competing interests

The authors have no known competing interests to declare.

## Ethical statement

The study complies with all relevant national and international ethical laws and guidelines. No ethical board recommendation was needed to work on *Cornu aspersum*.

